# Pex9 regulates TORC2 during the cellular response to oxidative stress

**DOI:** 10.64898/2026.01.25.701303

**Authors:** Mor Angel, Amir Szitenberg, Miri Carmi, Yeynit Asraf, Naama Barkai, Maya Schuldiner

**Affiliations:** Department of Molecular Genetics, Weizmann Institute of Science, Israel; The Mantoux Bioinformatics institute of the Nancy and Stephen Grand Israel National Center for Personalized Medicine, Weizmann Institute of Science, Israel

## Abstract

Peroxisomes are known for their detoxification role in cellular reactive oxygen species (ROS) homeostasis, yet their regulatory roles in cellular recovery from oxidative stress remain unclear. Here, using yeast, we systematically screened a panel of fluorescently tagged peroxisome-related proteins and identified widespread stress-induced changes in their localization. Notably, we discovered that the peroxisomal import receptor, Pex9, rapidly enters the nucleus upon H_2_O_2_ exposure, independently of the canonical oxidative stress regulators-Yap1 and Skn7, suggesting a non-canonical signaling function. We further show that Pex9 nuclear entry is modulated by Avo1, an essential Tor complex 2 (TORC2) subunit. In turn, Pex9 downregulates the essential TOR complexes kinase, Tor2, at both the transcriptional and protein levels. Finally, we show that this Pex9-TORC2 regulatory loop during oxidative stress is facilitated by the cellular envelope stress response. Our findings uncover an unexpected peroxisome-TORC2 signaling axis and highlight the importance of organelles in shaping global cellular responses to stress.

## Introduction

Cells constantly need to adjust to metabolic, physiological and stress alterations. Regardless of whether stress is sensed in the cytosol, or from inside organelles, the typical stress response includes a “sense and respond” pathway which starts from the “sensor” of changing conditions, continues with the modulation of a key regulator of transcription or translation and ends with cellular rewiring to re-establish homeostasis (*1–4*). For example, the canonical stress response for oxidative stress in the yeast *Saccharomyces cerevisiae (*hereafter referred to as yeast) is modulated by Yap1 and Skn7. This response includes a sensing step mediated by Gpx3 that directly detects H_2_O_2_, this is followed by activation of Yap1, a key transcription factor, by modulating its entrance to the nucleus, which in turn, leads to a transcriptional response consequently allowing for adapting and relieving the stress. Yap1 (as well as other, yet undiscovered proteins) also regulates Skn7 through phosphorylation, contributing to the cellular recovery (*5–8*). However, this pathway is initiated by cytosolic sensing, and it remains unclear whether additional “sense and respond” pathways are driven by cellular organelles.

Peroxisomes, ubiquitous organelles in eukaryotic cells, were named in 1966, by De Duve and Baudhuin who discovered the presence of peroxides, H_2_O_2_-producing oxidases as well as the detoxifying catalase, in these organelles (*9*). Indeed, it was later shown that peroxisomes play a role in the oxidation of fatty acids, a process that yields Reactive Oxygen Species (ROS) and heat (*10–12*). Since then, many studies have been published about the relationship of peroxisomes and oxidative stress. For example, it was shown that H_2_O_2_ can impact peroxisomal autophagy (i.e., pexophagy) (*13*, *14*), presumably to limit intrinsic ROS formation during times of oxidative stress. Later, it was demonstrated that oxidative stress leads to cytosolic accumulation of the peroxisomal H_2_O_2_ degrading enzyme, catalase, by blocking its peroxisomal import (*15*, *16*) suggesting a role in cytosolic detoxification. Furthermore, it was recently shown that peroxisomes aid in detoxifying mitochondrial ROS (*17*). Thus, the connection between peroxisomes and active detoxification of ROS during oxidative stress is inherent. However, it is still unclear whether their role extends beyond direct ROS elimination, to sensing and regulating the cellular response to, or recovery from, oxidative stress.

In this study, we set out to discover what is the active regulatory role of peroxisomal proteins in the cellular response to oxidative stress. We screened ∼150 fluorescently tagged peroxisome-related proteins in yeast and found that many of them change their cellular localization under oxidative stress. Among them, we observed that the peroxisomal import factor, Pex9 (*18*, *19*), immediately enters the nucleus in H_2_O_2_ treated cells, independently of the canonical oxidative stress response mediated by Yap1 and Skn7 (*20*), suggesting that it might carry a role as a non-canonical transcription modulator. Surprisingly, we found that the nuclear entry of Pex9 is modulated by Avo1, an essential subunit of the master regulator of cellular growth and metabolism, Tor complex 2 (TORC2) (*21*). Using RNA sequencing we show that Pex9 regulates an extensive set of transcripts amongst them the essential kinase of the TOR complexes, Tor2 (*22*, *23*). We show that Pex9 regulates Tor2 both at the RNA and protein levels. Finally, we demonstrate that this Pex9-TORC2 regulatory loop during oxidative stress, is facilitated by the cellular envelope stress response. Our data reveal that peroxisomal proteins have regulatory roles in cellular recovery from oxidative stress and uncover a novel signaling axis.

## Results

### Oxidative stress impacts the localization of peroxisome-related proteins

Peroxisomes have long been recognized for their role in ROS formation and detoxification. However, the mechanisms by which peroxisomal proteins manage and promote cellular response to, and recovery from, oxidative stress remain unclear. To get a hint of which proteins may regulate the transfer of information related to peroxisomes during oxidative stress, we first looked for striking changes in localization of peroxisomal, and peroxisomal-related, proteins, in the presence or absence of oxidative stress. To visualize the range of peroxisomal proteins in an unbiased manner, we imaged ∼150 proteins fused to a fluorophore at either their C or N-terminus (this was important to ensure the correct targeting of peroxisomal proteins that rely on their extreme C-terminus for peroxisomal import). We imaged cells following addition of hydrogen peroxide (H_2_O_2_) to induce oxidative stress (**Figure 1A**) and found several proteins that manifested changes in their localization or abundance (Figure **1B****, C, Supplementary Table 1**).

**Figure 1:**
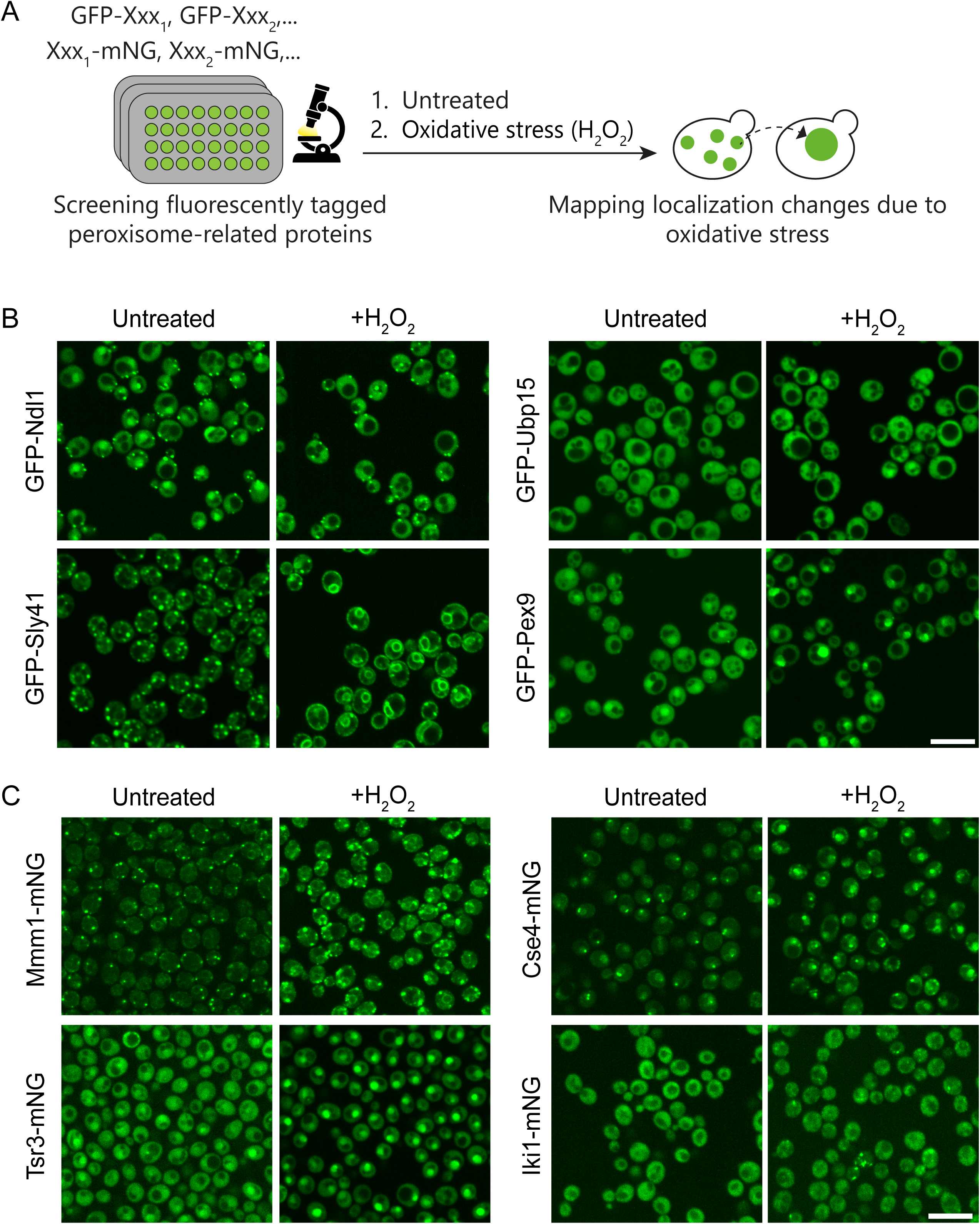
Peroxisome related proteins change their cellular localization in response to oxidative stress. (A) scheme describing the methodology for finding novel peroxisome-related stress response transcription modulators. Representative images of cells expressing fluorescently tagged peroxisome-related proteins on either their N-terminus (B) or C-terminus (C). Cells were imaged with and without H_2_O_2_ (2mM, 1 hr), and changes in their sub-cellular localization were observed. Scale bars=10µm.

The peroxisome-related protein collection included control strains for other cellular components (*24*), and surprisingly, one of these control strains, Sly41, had a striking change in cellular localization from punctuate pattern to ER, when H_2_O_2_ was applied. Sly41 is a protein involved in ER to Golgi trafficking (*25*, *26*). This suggests that oxidative stress either impacts its transport from ER to Golgi or else alters the rate of ER exit more globally-this remains to be followed up on.

Ubp15, Pex9 and Tsr3 entered the nucleus in oxidative stress, while Ndl1 lost its nuclear localization following H_2_O_2_ treatment. These striking movements suggest a regulatory role for these proteins during stress and prompted us to further investigate this avenue.

### Pex9 is a potential transcription modulator functioning under oxidative stress

In many cases, an integral part of the cellular response to stress is modulated by altering the transcriptome, that, in turn, allows cellular adaptation and recovery (**Figure 2A**). Therefore, to start exploring whether peroxisomal proteins contribute to cellular adaptation to stress by modulating the transcriptome, we focused on proteins that enter the nucleus under oxidative stress. Specifically, we chose to proceed with the peroxisomal protein Pex9 that exhibited dramatic nuclear entry in these conditions (**Figure 1B**). Pex9 is a central modulator of peroxisomal metabolism since it enable*s* the import of specific enzymes required for shifting the glyoxylate cycle into peroxisomes during fatty acid degradation (*18*, *19*, *27*). As such it enters peroxisomes during increased ROS production when yeast grow on oleate as a sole carbon source. However, it was not known that it can localize to the nucleus and its contribution to oxidative stress adaptation was never explored.

**Figure 2:**
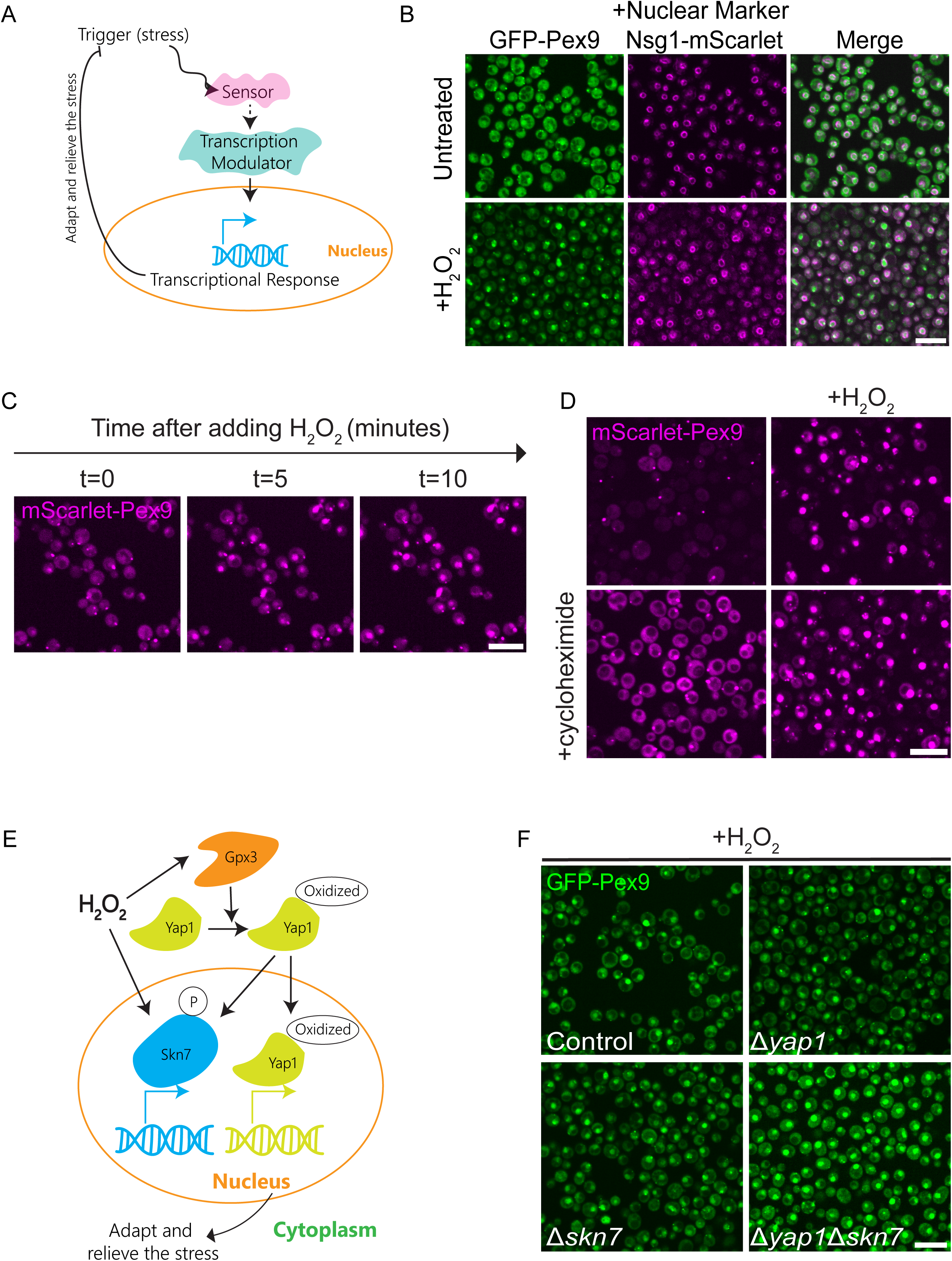
Pex9 rapidly enters the nucleus following oxidative stress, independently of the canonical oxidative stress response. (A) A scheme describing a typical stress response pathway. A sensor can detect stress, leading to a cascade of events culminating in a transcription modulator entering the nucleus to allow transcriptional adaptation. (B) Pex9 enters the nucleus under oxidative stress. GFP-Pex9 tagged cells were treated with H_2_O_2_ (2mM, 1 hr), and Nsg1 tagged with mScarlet was used to assess nuclear localization. Scale bars=10µm. (C) Pex9 enters the nucleus rapidly. mScarlet-Pex9 tagged strains were imaged before applying oxidative stress, and then every 5 minutes following H_2_O_2_ (2mM, 1 hr) treatment. (D) Newly synthesized proteins are not required for Pex9 response to oxidative stress. mScarlet-Pex9 cells were treated either with cycloheximide (µ200g/ml, 1 hr) and/or H_2_O_2_ (2mM, 1 hr). (E) A scheme illustrating the canonical oxidative stress response in yeast. Yap1 oxidation is mediated by Gpx3. Oxidized Yap1, together with the phosphorylated Skn7, yields a transcriptional response helping the cells to return to homeostasis following stress initiation. (F) The nuclear translocation of Pex9 is not modulated by the canonical stress response pathway. GFP-Pex9 tagged cells still display entrance to the nucleus after H_2_O_2_ (2mM, 1 hr) treatment also in the background of Δ*yap1*, Δ*skn7* or Δ*yap1*Δ*skn7*.

We verified that Pex9 enters the nucleus by tagging the nuclear periphery protein Nsg1 (*28*) (**Figure 2B**). Strikingly, the entry of Pex9 to the nucleus starts as early as 5 minutes after oxidative stress is applied, pointing to a “first line of defense” function in cellular recovery to oxidative stress (**Figure 2C**). In support of this, translation inhibition by cycloheximide did not prevent the entry of Pex9 to the nucleus, supporting the idea that the existing pool of Pex9 shifts its localization (**Figure 2D**). Interestingly, all four cargos that are known to be imported into peroxisomes by Pex9 (Mls1, Dal7, Fsh3 and Gto1) (*27*) are not shifted into the nucleus during oxidative stress, suggesting that entry into the nucleus of Pex9 represents an additional, non-cargo related, function (**Supplementary Figure S1A**). Furthermore, the main cargo adaptor, Pex5, that is a Pex9 paralog and highly similar structurally and functionally (*29*) does not enter the nucleus in response to H_2_O_2_ (**Supplementary Figure S1B**). These results emphasize that this phenomenon is not common among all peroxisomal cargo import proteins and represents a unique role of Pex9.

We then assayed whether the Pex9 shift in localization is impacted by the canonical stress response mechanisms to oxidative stress mediated by the transcription factors Yap1 and Skn7 (*20*). (**Figure 2E**). We found that deletion of *YAP1*, *SKN7*, or both did not affect the nuclear localization of Pex9 under oxidative stress (**Figure 2F**). Neither did deletion of genes encoding for the homologs of Skn7 and Yap1-Hms2 and Cad1 (*30*) (**Supplementary Figure S1C**). Altogether, the data suggest that Pex9 plays a role in an immediate cellular recovery mechanism induced by oxidative stress, that is independent of the Yap1/Skn7 axis.

### H_2_O_2_ induces global transcriptomic changes impacting a major route of peroxisomal protein import

Since Pex9 enters the nucleus immediately following oxidative stress, and this is not regulated directly by Skn7 and Yap1, we hypothesized that it elicits a unique transcriptomic signature. To determine if this is the case, we analyzed a control strain (same genetic background) as well as deletions of *PEX9*, *YAP1* and *SKN7*, before and after H_2_O_2_ treatment, looking for differential transcript levels.

Principal component analysis (PCA) shows that all H_2_O_2_ treated samples obtained a different transcriptional profile compared to the untreated samples, verifying that H_2_O_2_ has a general effect on the transcriptomic profile in all strains (**Supplementary Figure S2A**). As validation for the analysis, transcriptomic changes of the control strain clearly demonstrate that the addition of H_2_O_2_ worked as described previously in the literature. For example, genes that are well known to be upregulated via Yap1 due to H_2_O_2_ treatment such as *SOD1*, *SOD2* and *TSA1* (*20*, *31*, *32*) were indeed upregulated in the control strain, yet not in the Δ*yap1* strain (**Figures 3A-B**). To further verify our transcriptomic data using an orthogonal approach, we followed six randomly picked proteins, whose transcripts increased significantly and dramatically, for their abundance. Indeed, all proteins (C-terminally tagged, leaving their native promoters intact) showed higher signal upon H_2_O_2_ treatment (**Figure 3C**).

**Figure 3:**
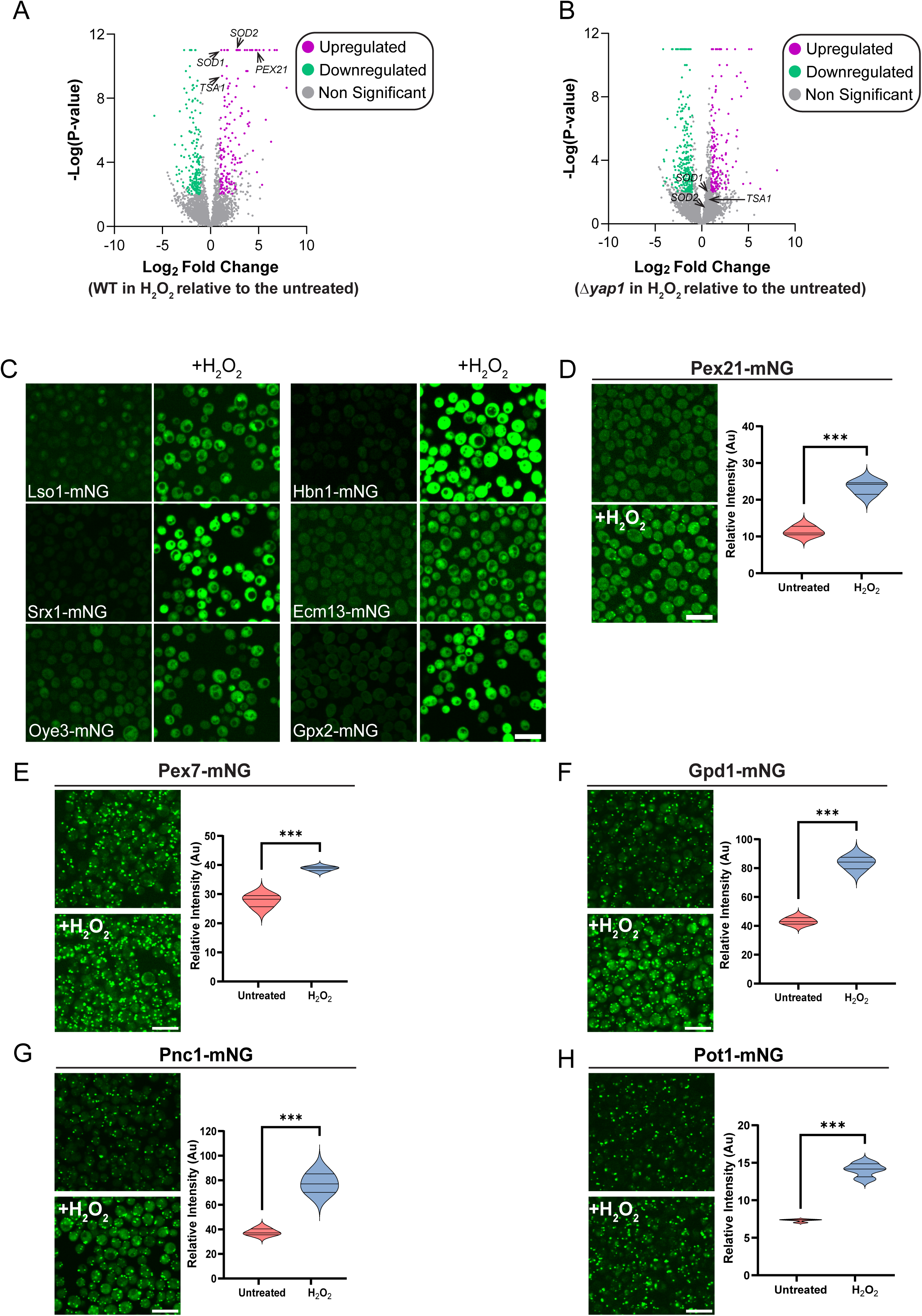
Transcriptomics reveal an increase in PTS2 proteins during oxidative stress. (A-B) Volcano plot showing genes that are upregulated (magenta), downregulated (green) or not significantly changed (gray) in cells treated with H_2_O_2_ (2mM, 1 hr) compared to untreated cells in (A) the control strain or (B) in the Δ*yap1* strain. Significancy was set at \*\*\**p-value*<0.001, log_2_fold change≥1. (C) Genes whose transcripts are upregulated in the control strain due to H_2_O_2_ treatment are also upregulated at the protein level. Lso1, Srx1, Oye3, Hbn1, Ecm13 or Gpx2 C-terminally tagged with mNeonGreen (mNG), thus preserving their native promoter regulation, were treated with H_2_O_2_ (2mM, 2.5 hr) and imaged at the same time and conditions as the untreated cells. Scale bars=10µm. (D-H) PTS2 machinery and cargos show increased expression level in H_2_O_2_ treated cells. Pex21, Pex7, Gpd1, Pnc1 or Pot1 C-terminally tagged with mNG were treated with H_2_O_2_ (2mM, 2.5 hr) and imaged at the same time and conditions as the untreated cells. Intensity was quantified and normalized to non-fluorescently tagged strains. Statistical significance was determined by *t*-Test, \*\*\**p*<0.001.

Notably, one transcript that increased dramatically due to oxidative stress was *PEX21* (**Figure 3A**). This change was also observed at the protein level (**Figure 3D**). Pex21 is a co-factor for the cargo adaptor Pex7 during the import of proteins that have a unique Peroxisomal Targeting Signal (PTS2) into peroxisomes. In support of, this we find that the abundance of PTS2 cargo adaptor Pex7 (*33*), also increased in H_2_O_2_ treated cells (**Figure 3E**). This increase clearly impacts their joint cargo, Gpd1, Pnc1 and Pot1 (*34–36*) (**Figure 3F-H**). Our results go hand in hand with previous findings demonstrating that the PTS2 proteins Gpd1 and Pnc1 help the cell to cope with osmotic pressure and that their levels in the cytoplasm and peroxisome are increased during stress (*37*). Overall, our data demonstrate that PTS2 machinery and cargos are impacted by H_2_O_2_ treatment, suggesting that they play an important role in coping with, or reacting to, oxidative stress.

### Pex9 downregulates Tor2 in response to oxidative stress

To reveal novel pathways for cellular recovery from oxidative stress regulated by Pex9, we analyzed the transcriptomic data by filtering for genes that were significantly changed in control cells exposed to H_2_O_2_, yet were unchanged in Δ*pex9* stressed cells. As suggested by its nuclear entry, we indeed detected 48 genes that significantly were up- or down-regulated in a Pex9-dependent manner in oxidative stress (**Figure 4A**).

**Figure 4:**
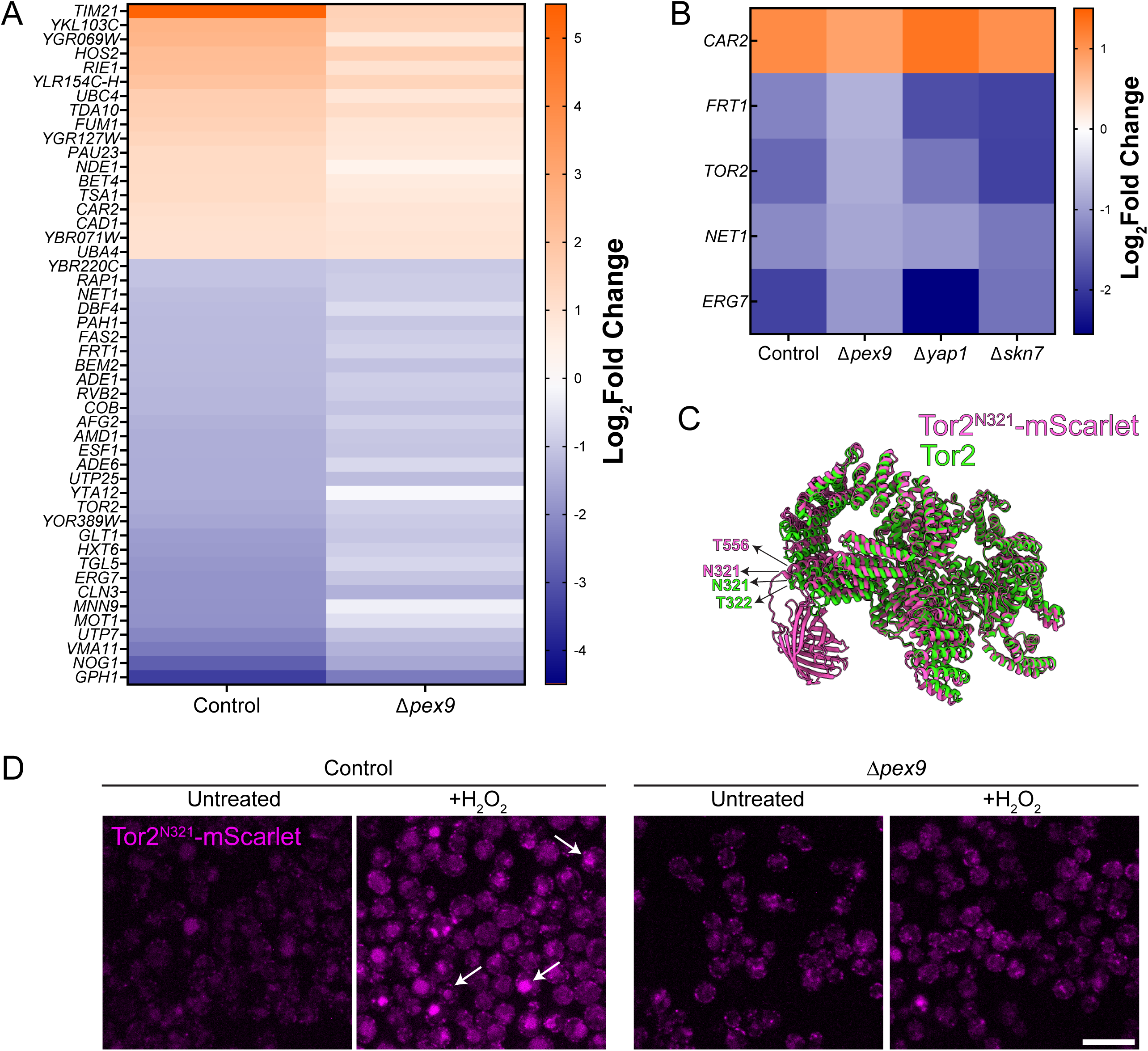
Pex9 downregulates TORC2 at both the RNA and protein levels. (A) Heat map showing the genes that are regulated by Pex9 in oxidative stress. Representation of the significantly changed genes in the control strain treated with H_2_O_2_ (2mM, 1 hr) and are not-significantly changed in Δ*pex9* cells. Significance was set at \*\*\**p-value*<0.001, log_2_fold change≥1. (B) Genes that are uniquely regulated by Pex9 (Changed significantly in all strains but not in Δ*pex9* cells following H_2_O_2_ (2mM, 1 hr) treatment). Significance was set at \*\*\**p-value*<0.001, log_2_fold change≥1. (C) AlphaFold model of Tor2 (green) and Tor2 tagged with mScarlet (magenta) at the N321 position, demonstrating that the fluorophore does not interfere with its structure. Position N321, where the scarlet was added, and the consequent amino acid (T) are shown on both models (position 322 or position 566). (D) Tor2 enters the vacuole during oxidative stress, and this is facilitated by Pex9. Tor2^N321^-mScarlet (control and Δ*pex9*) were treated with H_2_O_2_ (2mM, 1.5 hr). White arrows point to the vacuoles. Scale bar=10µm.

To focus on the genes that are regulated specifically and exclusively by Pex9 in oxidative stress we then filtered out those transcripts that were also affected by Δ*yap1* and Δ*skn7* strains. While many genes were shared with Yap1 and Skn7, suggesting that they may collaborate with Pex9 for some of their regulatory functions, this analysis revealed that Pex9 also impacts an exclusive list of transcripts, including the downregulation of the essential ergosterol biosynthesis enzyme, Erg7 (*38*) (also verified by protein levels (**Supplementary Figure S2B**)) and Tor2 (**Figure 4B**).

Tor2 is an essential subunit of both TOR complexes (TORC1 and TORC2) and is one of the most central metabolic regulators in the cell (*21*, *39*, *40*). We therefore evaluated whether the Pex9-mediated decrease in transcript levels impacts Tor2 also at the protein level. Since no Tor2-specific antibodies exist and fusing fluorophores to Tor2 at either terminus leads to its functional instability, we tagged Tor2 internally using CRISPR with the fluorophore mScarlet. We inserted mScarlet in position N321, that was previously shown to be compatible with its function (*41*) and modelled by AlphaFold (*42*) to ensure that insertion of this tag does not interrupt its fold (**Figure 4C**) or complex assembly (**Supplementary Figure S2C**). Deleting *PEX9* in the Tor2^N321^-mScarlet strain showed a clear effect (**Figure 4**). First, in the untreated cells the Tor2^N321^-mScarlet signal was lower in the control compared to the Δ*pex9* cells. Furthermore, during stress, the majority of the signal was detected within vacuoles, visualized by the vacuolar marker Vph1-mNG (*43*) (**Supplementary Figure S2D**), suggesting active degradation of Tor2. However, this was barely observed in the absence of Pex9 (**Figure 4D**), despite no general effect on vacuoles in this background (**Supplementary Figure S2E**).

Overall, these results show that Pex9 downregulates Tor2 both at the RNA and protein levels and suggests that it impacts signaling by TOR complexes under oxidative stress.

### Avo1, a TORC2 essential subunit, modulates the entry of Pex9 to the nucleus during oxidative stress

Our data show that Pex9 enters the nucleus rapidly when oxidative stress is applied, and this, in turn, could impact the activity of the TOR complexes (**Figure 2C**, **Figure 4B**). We therefore set out to uncover who transmits the stress signal to Pex9 thereby modulating its localization shift. To this end, we integrated a fluorescently tagged Pex9 into a whole genome deletion (*44*) or knockdown collection for essential genes (i.e., Decreased Abundance by mRNA Perturbation-DAmP) collection (*45*), looking for a gene whose absence affects the nuclear shift of Pex9 during oxidative stress (**Figure 5A**). Overall, we found 21 genes affecting Pex9 localization (**Supplementary Table 2**), among them, the most striking ones were knockdown of Protein Kinase C (*PKC1*) that is essential for cell wall remodeling during stress (*46*, *47*), and the deletion of the ubiquitin hydrolase *DOA4* (*48*), both of which enhanced the nuclear localization signal of Pex9. On the other hand, deletion of a dubious open reading frame that partially overlaps the heat shock induced transcription factor, *HSF1* (*49*, *50*) and knockdown of *AVO1*, a TORC2 essential subunit (*21*), prevented entry of Pex9 to the nucleus. This suggests that upon oxidative stress they play a role in either sensing the stress or transmitting the information to modulate Pex9 nuclear localization (**Figure 5B**).

**Figure 5:**
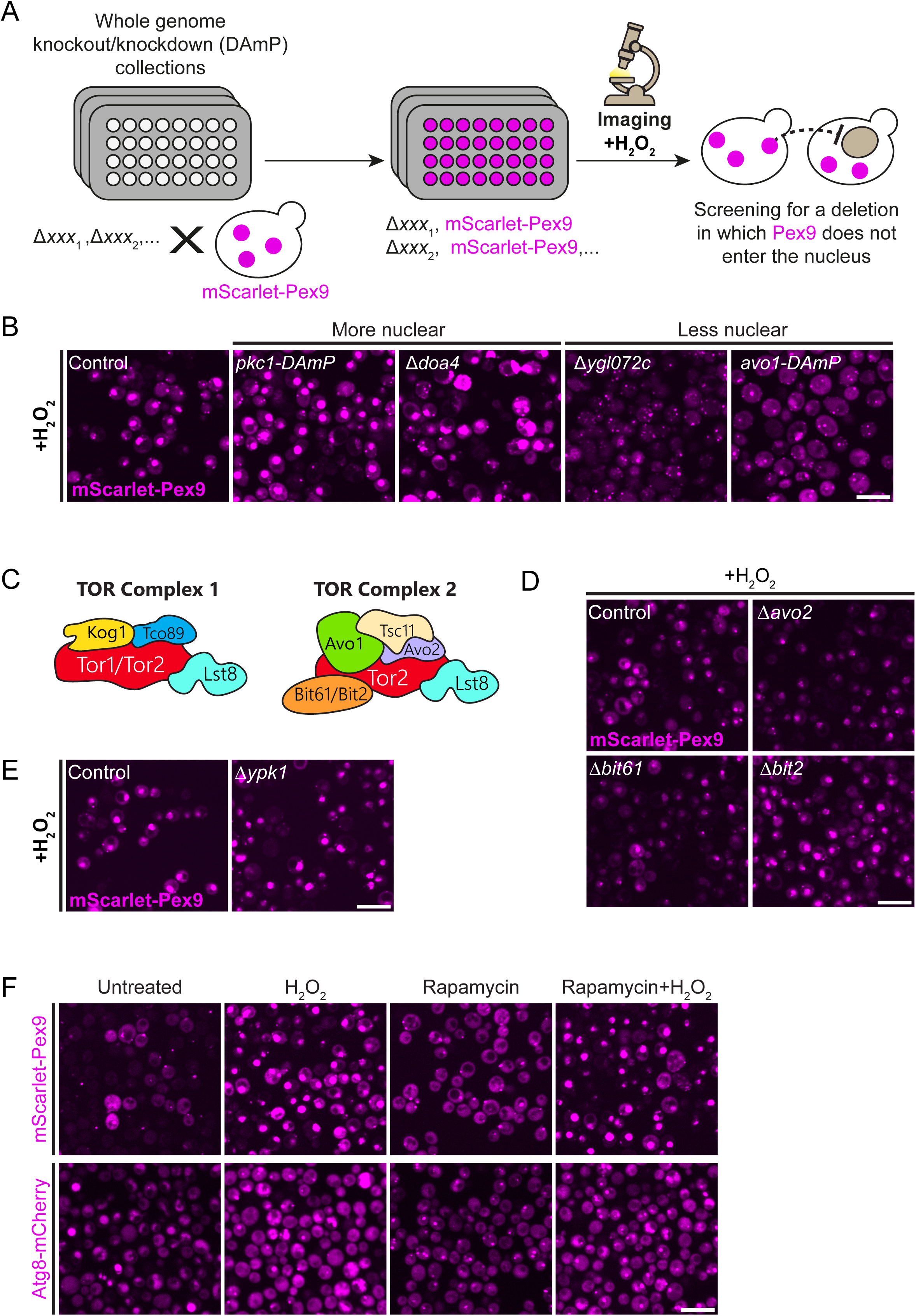
A systematic screen uncovers factors impacting Pex9 nuclear localization. (A) Scheme illustrating the process of finding candidates for modulating the nuclear entrance of Pex9. A whole genome deletion/knockdown (i.e., DAmP) was crossed with N-terminal fluorescently tagged Pex9 to get a combined collection. Then, strains were imaged in H_2_O_2_ (2mM, 1 hr) and nuclear localization of Pex9 was assessed. (B) The systematic screen that was described at (A), revealed that knocking down *PKC1* or deleting *DOA4* led to a stronger nuclear signal of Pex9, and deleting *YGL072C* or knocking down avo1 prevented the entrance of Pex9 to the nucleus in oxidative stress. Scale bars=10µm. (C) An illustration representing the components of TOR complex 1 (TORC1) and TOR complex 2 (TORC2). Tor2 can function in TORC1 or TORC2. (D) The non-essential components of TORC2 (i.e., Avo2, Bit61, Bit2) were deleted and Pex9 tagged in the N-terminus with mScarlet. Cells were imaged after H_2_O_2_ (2mM, 1 hr) treatment. (E) Deletion of TORC2 downstream kinase (*YPK1*) does not prevent the entrance of Pex9 to the nucleus under H_2_O_2_ (2mM, 1 hr) treatment (F) Inhibition of TORC1 by Rapamycin (200nM, 1 hr) does not prevent the entrance of Pex9 to the nucleus in H_2_O_2_ (2mM, 1 hr). Atg8-mCherry was used as a positive control to assess rapamycin treatment effectiveness.

Since we observed that Avo1, a component of TORC2, affects Pex9 localization, and since systematic screens suffer from false negatives, we manually assayed whether the other non-essential components of TORC2 (i.e., Avo2, Bit61, Bit2) (*51*) play a role in the nuclear localization of Pex9 under oxidative stress. We found that no other component impacted Pex9 localization to the same extent as *AVO1* deletion does (**Figure 5C-D**). We then assayed whether this activity was directly mediated by TORC2 or rather was a consequence of activation of its downstream kinases. Deleting *YPK1*, the kinase that is downstream to TORC2, or *SCH9*, the downstream kinase to TORC1 (*52*), did not prevent the entrance of Pex9 to the nucleus (**Figure 5E, Supplementary Figure S2F**), suggesting that this activity is directly modulated by TORC1/2 themselves. Since Tor2 can function as the kinase in both TORC1 and TORC2 (*53*) (**Figure 5C**) we deconvoluted which complex is important for this activity by using Rapamycin, an inhibitor that acts only on TORC1, but does not generally suppress the activity of TORC2 (*54*). Atg8, which was shown to change its cellular localization due to Rapamycin (*55*), was clearly impacted by the treatment, showing that TORC1 was indeed inhibited. Yet, Rapamycin did not diminish the nuclear entry of Pex9, suggesting lack of contribution from TORC1 to Pex9 localization changes in these conditions **(Figure 5F)**. Our data therefore support a direct role for TORC2 in Pex9 regulation following oxidative damage.

### Osmotic stress downregulates Tor2 and promotes Pex9 nuclear entry as part of negative feedback signaling loop

TORC2 is known for its role in plasma membrane (PM) maintenance and plays a role in coping with cellular envelope stress, that could be triggered by many stress types including oxidative stress (*56*, *57*). Specifically, it was shown that Avo1 contains pleckstrin homology (PH) domains that are specific for binding phosphatidylinositol-4,5-*bis*phosphate (PI4,5P2) and a conserved region in the middle (CRIM) domain that is required for the activation of Ypk1 which phosphorylates targets that control PM homeostasis (*58*, *59*). Therefore, we asked if reducing cellular envelope stress by applying sorbitol, an osmolyte that decreases PM tension (*60*), prior to H_2_O_2_ treatment, would affect Pex9 nuclear localization. Remarkably, we observed that sorbitol led to a milder nuclear entry of Pex9 (**Figure 6A**). This suggests that the sensing of Avo1, which ultimately sends Pex9 to the nucleus, might be mediated by the indirect impact of oxidative stress on cellular envelope tension.

**Figure 6:**
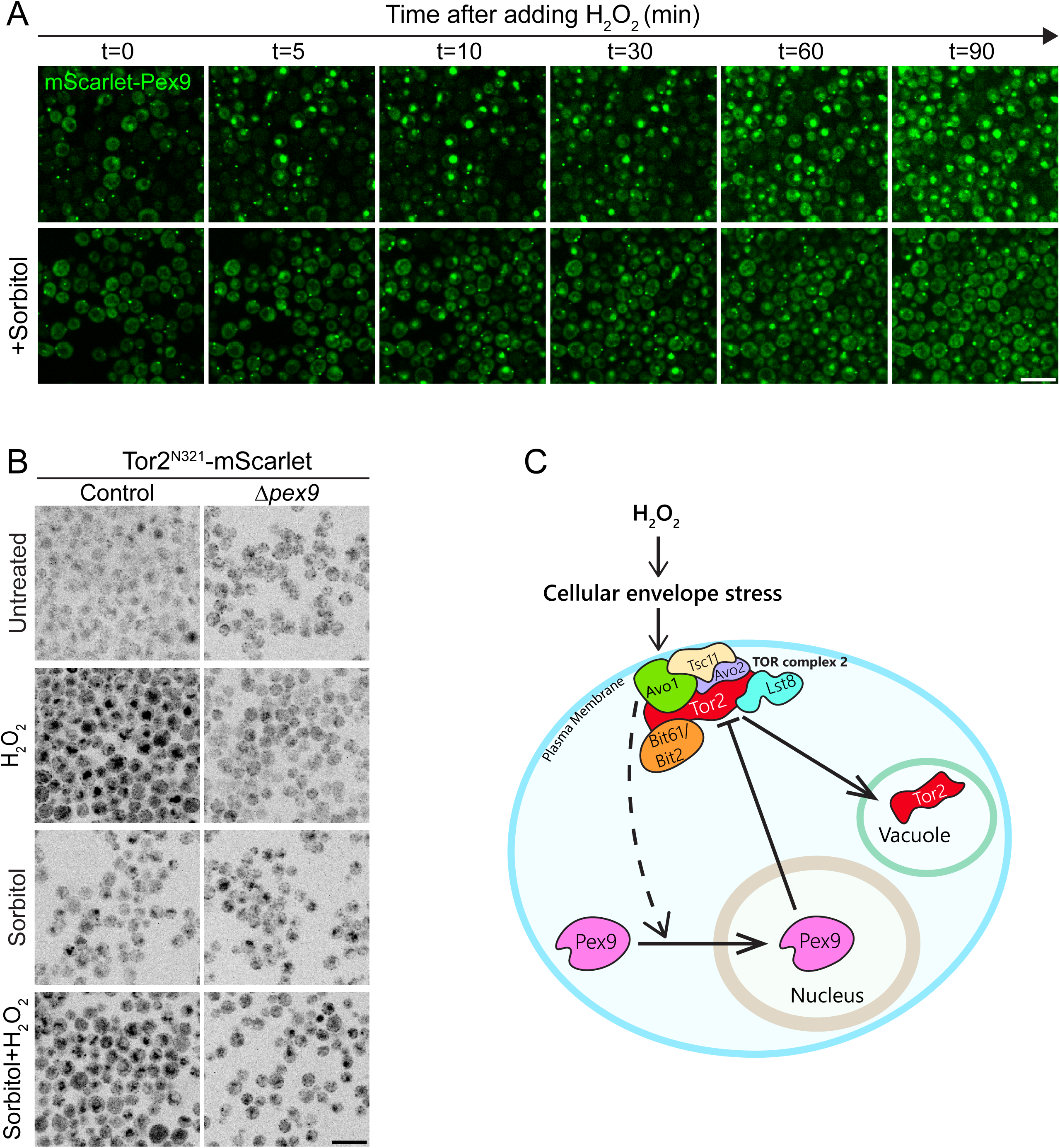
Pex9 creates a feedback loop with TORC2 modulated by osmotic stress. (A) mScarlet-Pex9 tagged cells were treated with H_2_O_2_ (2mM, 1 hr) to induce oxidative stress, sorbitol (1M, 4 hr) to decrease PM tension, or both, and images were taken every 5 minutes from initiation of oxidative stress. All images were taken in the same exposure time and intensity, yet cells that were pre-treated with sorbitol showed higher expression level due to osmotic stress, therefore the histogram of brightness/contrast was adjusted to all sorbitol treated cells, in comparison to all non-sorbitol treated cells. Scale bars= 10µm. (B) Tor2 enters the vacuole in osmotic stress, and this is independent of Pex9. Tor2^N321^-mScarlet cells (control and Δ*pex9*) were treated with sorbitol (1M, 4.5 hr overall) and/ or H_2_O_2_ (2mM, 1.5 hr), (C) A suggested model showing the feedback regulatory loop of Pex9 and TORC2 in oxidative stress. Oxidative stress (induced by H_2_O_2_ treatment) leads to cellular envelope stress, that can be sensed by TORC2 subunit Avo1. Avo1 modulates the entrance of Pex9 to the nucleus, which, in turn, downregulates Tor2 in the mRNA and protein level, therefore down regulating the TORC2 activity. Tor2 is directed to the vacuole either at oxidative stress or osmotic stress.

Since it was demonstrated that TORC2 is downregulated in cellular envelope stress (*60*, *61*), we asked whether adding sorbitol would affect not only Pex9 cellular localization in oxidative stress, but also Tor2. Surprisingly, we found that Tor2 is localized to the vacuole in sorbitol-treated cells, regardless of their oxidative state and independently of Pex9 presence (**Figure 6B**). This suggests that osmotic stress leads to downregulation of Tor2, yet this process is not modulated solely by Pex9. In addition, it implies that the downregulation of Tor2 by Pex9 is just one of many ways that enable Tor2 regulation.

Altogether, we suggest that Pex9 plays an important role in coping with cellular oxidative stress, and that this process is independent of the canonical oxidative stress pathways regulated by Yap1 and Skn7. We speculate that once oxidative stress ensues, Avo1 senses the impact on the cellular envelope, modulating the rapid entrance of Pex9 into the nucleus, which impacts the RNA levels of many proteins including down regulation of Tor2 (**Figure 6C**). This negative feedback loop reveals a novel stress response adaptation mechanism for cells to deal with oxidative stress and demonstrates for the first time that peroxisomal proteins have an active regulatory role in maintaining cellular homeostasis under stress.

## Discussion

In this study we showed that a peroxisomal protein holds a regulatory role in dealing with cellular oxidative stress. By screening the repertoire of peroxisome-related proteins, we found that many of them changed their localization due to oxidative stress (**Figure 1**). Specifically, we were interested in further examining proteins that enter/exit the nucleus thus having a potential capacity to impact RNA levels. Although Ubp15, Ndl1 and Tsr3 changed their nuclear localization due to oxidative stress (**Figure 1B-C**) and each should merit their own investigation, we chose to proceed with Pex9 for several reasons. First, it was not known to have a role in cellular adaptation to oxidative stress; second, it displays a striking change in nuclear localization following H_2_O_2_ treatment; and finally, it is a central peroxisomal protein. We show that Pex9 enters the nucleus rapidly after oxidative stress is applied, independent of protein synthesis (**Figure 2C-D**), emphasizing that Pex9 participates in an immediate cellular adaptation to stress. Importantly, the localization of Pex9 to the nucleus is not modulated by the canonical oxidative stress transcription modulators-Yap1 and Skn7 (**Figure 2F**), implying that Pex9 plays a role in a non-canonical stress response pathway.

The transcriptomic analysis reveals a set of genes that are regulated exclusively by Pex9 in these conditions. Although we verified that these genes are indeed regulated by Pex9 at the protein level (**Supplementary Figure 2B, Figure 4D**), it is still unclear if Pex9 regulates these genes directly through a transcriptional role or through an indirect nuclear role on mRNA stability or export. Interestingly, Pex9 contains a tetratricopeptide repeat domain, highly similar to a domain in Tfc4, a subunit of RNA polymerase III transcription initiation factor complex (*29*, *62*), proposing that Pex9 might not serve as a transcription factor by itself, yet could be modulating transcription elongation.

One of the genes that was downregulated by Pex9 in oxidative stress is *TOR2*. Tor2 is an essential protein and is the kinase of both TOR complexes-TORC1 and TORC2. Its homolog, Tor1 can only assemble into TORC1 making it non-essential unlike Tor2 (**Figure 5C**). It is poorly understood in which cases Tor2 replaces Tor1 in TORC1, yet it is known that when Tor1 is absent, Tor2 can compensate (*63*). Here, by using CRISPR, we visualize Tor2 with a strong fluorophore (**Figure 4C-D**). This might enable to further investigate the cellular localization of Tor2 under various conditions, and therefore shed light on this enigmatic protein (*41*).

We find that Avo1, a subunit of the TORC2, modulates the entry of Pex9 to the nucleus. It was previously described that TORC2 plays a role in PM and cell wall homeostasis, therefore detecting cellular envelope stress (*56*, *57*). Specifically, Avo1 is essential for the activation of the protein kinases Ypk1 and/or Ypk2, which phosphorylate targets regulating PM homeostasis (*58*). Since the deletion of *YPK1* does not affect Pex9 nuclear localization (**Figure 5E**), we suspect that Pex9 entrance to the nucleus is modulated by Avo1 through a different pathway, and not via the canonical Ypk1 mechanism.

Importantly, there is emerging evidence that ROS can impact TORC2 activity. For instance, it was demonstrated that TORC2-Ypk1 signaling restrains ROS accumulation, and that TORC2 regulates actin polarization by both vacuole-related and mitochondria-mediated ROS (*64*, *65*). Furthermore, Pkc1, that can be phosphorylated by TORC2, is part of the cell wall integrity (CWI) pathway, which plays a role in cell wall remodeling during stress (*46*, *47*). Recently it was demonstrated that the CWI pathway also negatively regulates TORC2-Ypk1/2 signaling during stress (*61*). Our data demonstrate that knocking down *PKC1* enhanced Pex9 nuclear localization (**Figure 5B**). Therefore, it is possible that Pex9 compensates for the absence of Pkc1, which in turn helps to deal with the cellular envelope stress.

While much work is required to unravel the entire interplay of TOR complexes and peroxisomes it is becoming clear that this interplay is important and conserved. For example, it was recently shown that in mammalian cells the tuberous sclerosis complex (TSC) and the ataxia-telangiectasia mutated (ATM) proteins are localized to peroxisomes by PEX5, and this suppresses mTORC1 (i.e., the mammalian homolog of yeast TORC1) in response to ROS and induces autophagy (*13*, *66*). It was also demonstrated that peroxisomal import stress activates the integrated stress response and downregulates the mTORC1 pathway (*67*). In yeast, abiotic stress leads to increased *de novo* biogenesis of peroxisomes, mediated by the inhibition of TOR signaling and activation of heat shock signaling (*68*). However, all these findings are focused on TOR complex 1, and we still lack information about TOR complex 2 in relation to peroxisomes.

Finally, it was recently suggested that TORC2 participates in a feedback loop to control active sterol levels at the PM as a general mechanism to adapt PM stresses, such as heat shock and hyperosmotic shock (*69*). We also find connections to ergosterol biosynthesis (regulation of Erg7 and Hsf1 controlling Pex9 nuclear entry). Indeed, we found that pre-treating oxidative-stressed cells with sorbitol, that was previously shown to decrease PM tension and leads to TORC2 inactivation (*60*), restricted Pex9 entrance to the nucleus in oxidative stress (**Figure 6A**). This suggests that Avo1 modulates the nuclear localization of Pex9 by sensing the cellular envelope stress. By observing Tor2, we uncover that osmotic stress leads to Tor2 vacuolar localization, in the absence or presence of Pex9, suggesting that TORC2 downregulation in these conditions occurs through trafficking Tor2 to the vacuole, independently of Pex9 (**Figure 6B**).

More globally, previous reports alongside our findings showcase that TORC1/2, the most central and conserved complexes in stress response and metabolism signaling, are modulated in a peroxisome-related manner. This suggests that indeed cells integrate sensing also from organelles into their general stress response modules for eliciting an optimal stress response.

Altogether, we show that the peroxisomal protein Pex9 has an active regulatory role in modulating an immediate cellular response to oxidative stress, facilitated by cellular envelope tension, and creating a feedback regulatory loop with TORC2 (**Figure 6D**). This emphasizes the important role that peroxisomal proteins play in recovery from oxidative stress in a broad cellular context.

## Materials and Methods

### Yeast strains and plasmids

All yeast strains used in this study are listed in **Supplementary Table 3**. Strains were constructed either by transformation, using the lithium acetate-based transformation protocol (*70*) or were picked from yeast libraries and were validated by PCR. All plasmids used are listed in **Supplementary Table 3**. Primers for transformations (cassette and validation check) were designed with the Primers-4-Yeast web tool (https://www.weizmann.ac.il/Primers-4-Yeast/) (*71*). For the initial H_2_O_2_ screen that is described in figure 1, the mini-peroxisomal libraries were imaged (*72*). All hits from the screening of this library are listed (**Supplementary Table 1**).

### Yeast growth and treatments

Yeast cells were grown on solid media containing 2.2% agar (AGA0X; Formedium) or liquid media, at 30°C. For selections, the following antibiotics were applied according to the corresponding strain (see **Supplementary table 3**): nourseothricin (NAT, WERNER BioAgents) 0.2 gr/liter, geneticin (G418, Formedium) 0.5 gr/liter, Hygromicin (Hygro, Gold Biotechnology) 0.5gr/liter. Yeast cells were grown on YPD (YPD broth 5% Formedium #CCM02XX: 2% peptone, 1% yeast extract, and 2% glucose). For strains that can grow on media with no uracil, synthetic minimal media was used (SD; 0.67% yeast nitrogen base without amino acids and with ammonium sulfate #CYN04XX; Formedium, 2% glucose). SD media was supplemented with amino acid OMM mix (*73*) (i.e., SD AA complete) for back dilution and for imaging. For overnight cultures SD media without Uracil (Ura) was used.

For inducing oxidative stress, cells were grown for 3 hours to reach mid-log phase, then were treated with 2mM H_2_O_2_ (Bio-Labs, 085503) for an additional 1 hr. µ200g/ml Cycloheximide (Sigma, C4859) or 200nM rapamycin (Sigma, 553210) treatment was applied together with the H_2_O_2_, for a 1 hr treatment overall, and 1M D-Sorbitol (Sigma, S1876) was added to the cells before reaching mid-log, for 4 hr overall.

### Yeast library generation

The mScarlet-Pex9-full genome deletion library generation was performed as previously described (*74*). In short, a RoToR array pinning robot (Singer Instruments) was used to mate the donor strain with the parental deletion or DAmP library (*44*, *45*). (See more details in **Supplementary Table 3**).The mating was followed by sporulation and selection to generate the desired haploid a library (*74*, *75*). Finally, a combination of NATR, G418R and media without Ura was employed to select the strains that were successfully completed. All hits from the screening of this library are listed (**Supplementary Table 2**).

### RNA extraction

Cells were grown to mid-log in YPD, and 2mM H_2_O_2_ was added for 1 hr prior to cell wall digestion. Cells were washed with DDW, then RNA was extracted using the YeaStar RNA kit (Zymo, ZR-R1002) according to the manufacturers protocol, following DNAse I treatment (Zymo, ZR-E1010). RNA concentration and quality were assessed by nanodrop and TapeStation.

### Bulk MARS-Seq

RNA-seq libraries were prepared at the Crown Genomics Institute of the Nancy and Stephen Grand Israel National Center for Personalized Medicine, Weizmann Institute of Science. A bulk adaptation of the MARS-Seq protocol (*76*, *77*) was used to generate RNA-Seq libraries for expression profiling of the control strain, Δ*skn7*, Δ*yap1*, Δ*pex9*, treated and untreated in 3 biological and 3 technical replicates. Briefly, 30 ng of input RNA from each sample was barcoded during reverse transcription and pooled. Following Agencourt Ampure XP beads cleanup (Beckman Coulter), the pooled samples underwent second-strand synthesis and were linearly amplified by T7 *in vitro* transcription. The resulting RNA was fragmented and converted into a sequencing-ready library by tagging the samples with Illumina sequences during ligation, RT, and PCR. Libraries were quantified by Qubit and TapeStation as well as by qPCR for the *ACT1* gene as previously described (*76*, *77*). <u>Act1 F:</u> *TCATTGCTCCTCCAGAAAGAA;* <u>Act1 R:</u> *TGTGGTGAACGATAGATGG* Sequencing was done on a NovaSeqX plus using half a kit of 1.5B 100 cycles, allocating 800M reads in total (Illumina).

### RNA-seq Analysis

To quantify gene expression, we initially trimmed the sequence read data to remove Illumina adaptor sequences and poly-A sequences, as well as low quality read sections, using cutadapt (*78*). Resulting reads shorter than 30 bp were excluded from downstream analyses. Then the reads were mapped to the R64 reference genome assembly of *S. cerevisiae* S288 with STAR (*79*). To account for the yeast’s small introns, compared to STAR’s default expectations, we limited the accepted intron size to 0-5000 bp. A count table was prepared with HTSeq-count (*80*), only considering reads that map to the 3’ UTR regions, which was estimated to exists in the 1000 first 3’ bases of the transcript region, as indicated in the RefSeq annotations. Reads sharing a UMI sequence, which mapped to the same gene, where deduplicated to avoid technical effects. Differentially expressed genes were identified with DESeq2 (*81*), with the Benjamini and Hochberg procedure for multiple testing (*82*). Genes with p-value ≤ 0.01 and |log2-Fold-Change| ≥ 1 (i.e. at least two times count difference) were differentially expressed, as long has the sample with the highest count had at least 20 reads.

### Fluorescence microscopy

Cells were grown overnight at 30°C on the desired selection, then culture was back-diluted into 384-well plates in SD supplemented with amino acid OMM. 2mM H_2_O_2_ was added 1 hr prior imaging. After 4 hr, 50 μl from each well were transferred to a glass-bottom 384-well microscopy plate (Azenta Life Sciences) coated with concanavalin A (SigmaAldrich). The logarithmic phase cultures were allowed to bind to the bottom of the plate for 20 min. After incubation, wells were washed twice with SD complete media (with or without H_2_O_2_) to remove non-adherent cells. Plates were imaged using an Olympus IX83 microscope, 60x oil-immersion objective (NA = 1.42). VisiScope Confocal Cell Explorer system (Visitron Systems) is equipped with spinning disk scanning unit (CSU-W1, Yokogawa) and an Edge sCMOS camera (PCO). The system was operated using VisiView software (V3.2.0, Visitron Systems).

For high throughput screens, the plates were transferred to an Olympus automated inverted fluorescent microscope system using a robotic swap arm (Peak Robotics). Cells were imaged at room temperature (RT) in SD complete media using an Olympus UPLFLN 60X air lens (NA 0.9), a CSU-W1 Confocal Scanner Unit (Yokogawa), equipped with a 50 µm pinhole disk, and with an ORCA-flash 4.0 digital camera (Hamamatsu), using the ScanR software (V3.2.0). All the steps in the automated imaging process were done by an automated imaging platform EVO freedom liquid handler (TECAN).

### ScanR analysis and statistics

All experiments were carried out at least three biological replicates.

The intensity analysis of the PTS2 proteins was carried out using ScanR acquisition software (V 3.2.0). Yeast cells were segmented using neural networks-based approach applied to the brightfield images, and the mean fluorescence intensity was quantified from the 488 nm images for each cell. The Pot1-mNG strain, with and without H_2_O_2_, was imaged in different intensity than all other PTS2 strains, due to differences in expression level and saturation.

Then, the intensity values were normalized to a non-fluorescent strain, that was imaged at the same conditions and experiment. For each condition, intensity values were averaged across the three biological replicates.

All statistical analysis was conducted using GraphPad Prism 10.0. Statistical significance for the PTS2 proteins intensity was carried out using *t*-test. Heat maps and volcano plots were plotted using GraphPad Prism as well. To include differentially expressed genes in the volcano plots and the heat map, we required that *p*-value ≤0.01 the absolute value of log2Fold Change≥1, and that at least one sample had a count of ≥20 reads.

### Creating the Tor2^N321^-mScarlet strain using CRISPR

Gene editing of the Tor2^N321^-mScarlet strain was carried out using the bRA89 plasmid (*83*), which encodes Cas9, a target-specific guide RNA, hygromycin and ampicillin resistance. A 20-bp guide RNA targeting *TOR2* with a PAM at positions 992–994 bp was designed and cloned into the bRA89 plasmid between the BplI sites.

The repair fragment was amplified from the mScarlet plasmid using a forward primer containing homology to *TOR2* including the N321 region, a GGGGS linker, and the N-terminus of mScarlet. The reverse primer included the C-terminus of mScarlet and Tor2 homology downstream of N321, incorporating a PAM-disrupting mutation (TGG → TAG). The plasmid and repair fragment were introduced into yeast cells, and positive colonies were verified. All primers that were used are listed at **Supplementary Table 4**.

### Structural modeling of Tor2

Protein structure predictions were performed using AlphaFold 3 (*42*). The alignment of the Tor2^N321^-mScarlet and the untagged Tor2 model, as well as all visualizations were carried out using ChimeraX version 1.11 (UCSF). N-terminal residues (amino acids 1-74) were excluded from alignments due to low confidence score (pLDDT<40).

## Acknowledgments

The robotic system in the Schuldiner laboratory was purchased through the kind support of the Blythe Brenden-Mann Foundation. M.S. is Incumbent of the Dr Gilbert Omenn and Martha Darling Professorial Chair in Molecular Genetics. RNA-seq analysis was done with critical advice from Inbal Bolocan-Nachman at the Crown Genomics Institute of the Nancy and Stephen Grand Israel National Center for Personalized Medicine, Weizmann Institute of Science. We thank Maria-Del-Rosario Valenti for her help with visualizing the Tor2 models using ChimeraX. We thank Michael N. Hall (University of Basel) for sending yeast strains (VA102, VA43 and TB50a) (*41*) with tagged Tor2. We are grateful to Dr. Naama Zung and Noga Preminger for critical reading of the manuscript.

## Author contributions

Conceptualization: MA, MS; Methodology: MA, MC, NB, MS; Investigation: MA, MC; Supervision: NB, MS; Data analysis: MA, AS, YA; Writing-original draft: MA, MS; Writing-review & editing: All authors

## Data and materials availability

All relevant data can be found within the article and its supplementary materials. The raw and processed data of RNA sequencing in the current study were deposited on GEO (accession number GSE316518).

## Supplementary Figure legends

**Supplementary Figure S1:**
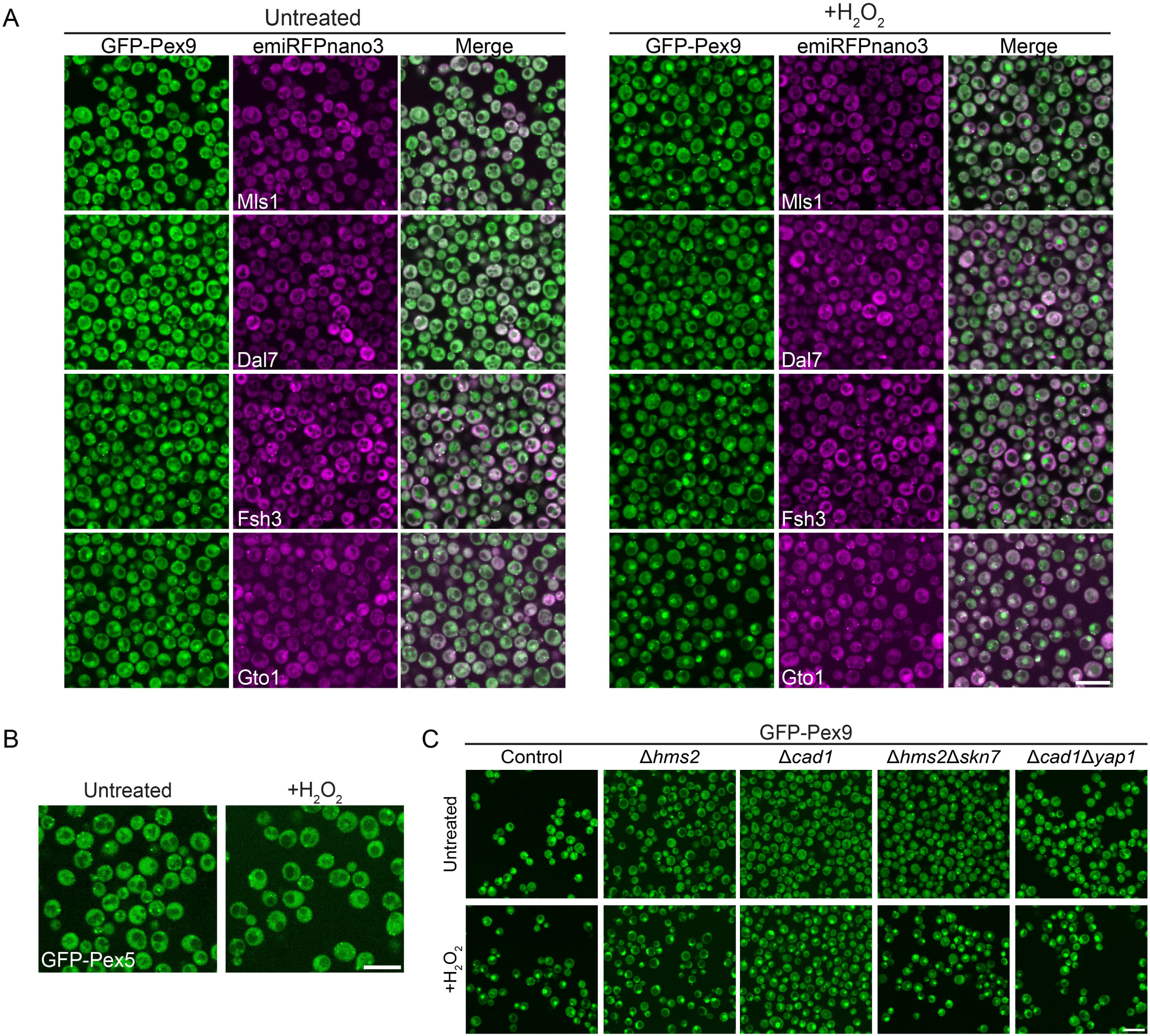
(A) Pex9 enters the nucleus in H_2_O_2_ (2mM, 1 hr) treated cells, yet its cargos (Mls1, Dal7, Fsh3 and Gto1) do not change their localization in these conditions. Pex9 cargos (magenta) were conjugated to emiRFPnano3 fluorophore at the C termini. Scale bars= 10µm. (B) Pex5 tagged with GFP in its N-terminus shows no change in cellular localization due to oxidative stress treatment H_2_O_2_ (2mM, 1 hr). (C) Deletion of Yap1 and Skn7 homologs -Cad1 and Hms2, does not change Pex9 nuclear localization in oxidative stress. GFP-Pex9 cells with the deleted genes were treated with H_2_O_2_ (2mM, 1 hr).

**Supplementary Figure S2:**
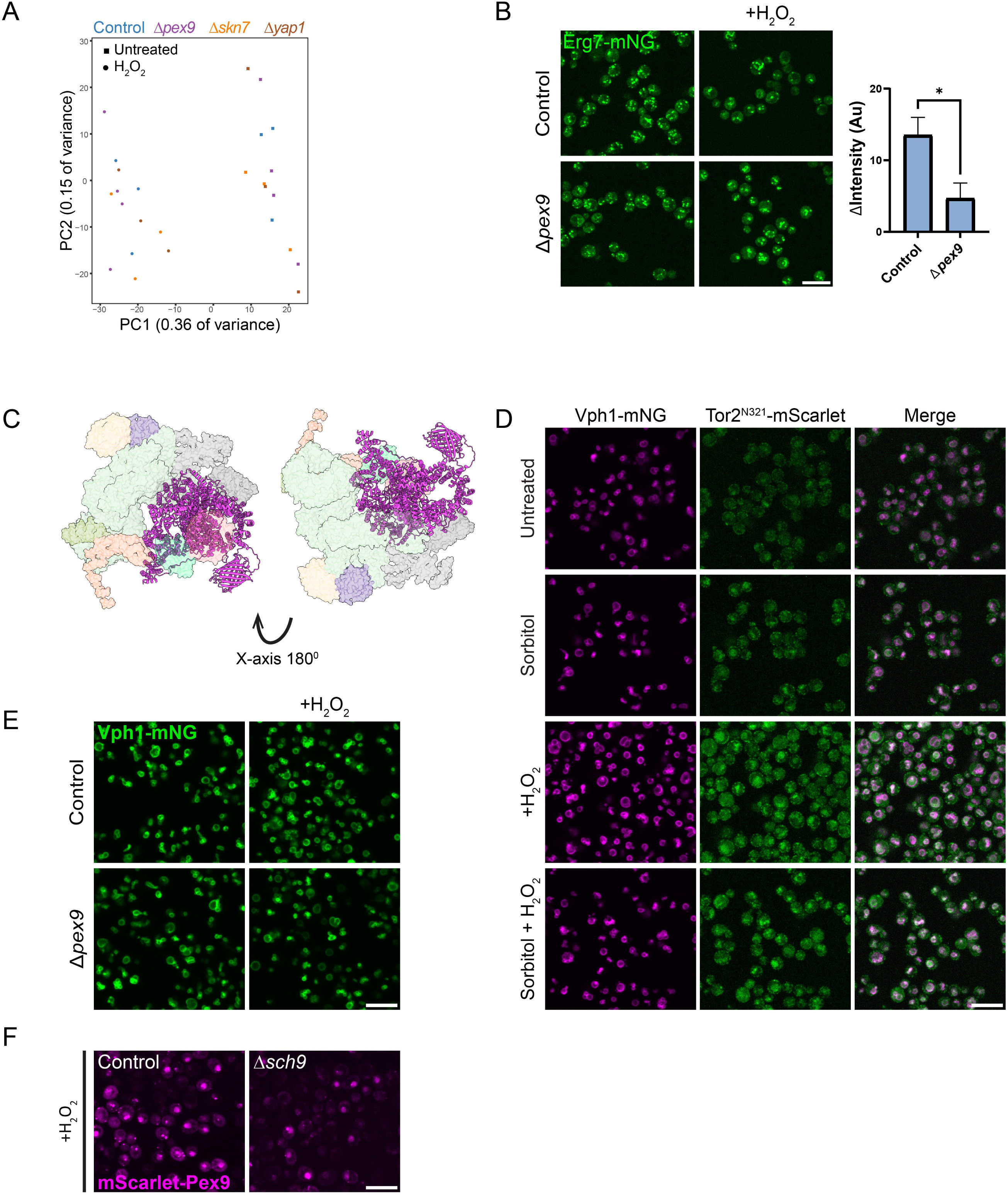
(A) PCA map showing the transcriptional profile of untreated and H_2_O_2_ (2mM, 1 hr) treated control, Δ*pex9*, Δ*skn7* and Δ*yap1* strains. (B) Erg7 was tagged with mNG in the C-terminus in control and Δ*pex9* cells to validate the transcriptomics data. Cells were treated with H_2_O_2_ (2mM, 1 hr), then the signal was quantified and the delta between the untreated and treated cells (ΔIntensity) was calculated. Statistical significance was determined by *t*-Test, \**p*<0.05 (C) The Tor2 complex that is shown was previously modeled using electron microscopy (Protein Data Bank accession code 6EMK) (*22*). Tor2^N321^-mScarlet was aligned using Chimera X, to ensure that the tagging of Tor2 does not interrupt with the complex structure. (D) Vph1 (magenta) was tagged with a fluorophore to validate that Tor2 (green) is sequestered within the vacuole in osmotic and oxidative stress. Cells were treated with H_2_O_2_ (2mM, 1.5 hr) and/or with sorbitol (1M, 4.5 hr). Scale bars= 10µm. (E) Control and Δ*pex9* Vph1-mNG tagged cells were treated with H_2_O_2_ (2mM, 1.5 hr) to ensure that deletion of *PEX9* does not change the vacuolar morphology. (F) Deletion of TORC1 downstream kinase (*SCH9*) does not prevent the entrance of Pex9 to the nucleus under H_2_O_2_ (2mM, 1 hr) treatment.

